# Alternative reproductive tactics and evolutionary rescue

**DOI:** 10.1101/2023.06.26.546260

**Authors:** Robert J Knell, Jonathan M. Parrett

## Abstract

Alternative reproductive tactics (ARTs), whereby males within a species exhibit qualitatively different behaviours in their pursuit of matings, are common throughout the animal kingdom. Here, using an individual-based modelling approach, we consider the possible impacts on adaptation and evolutionary rescue arising from one class of ARTs, those expressed through condition-dependent polyphenism such that high-quality, well-adapted males compete aggressively for mates and low-quality, poorly adapted males attempt to acquire matings via other, non-aggressive behaviours. When the ART is simultaneous, meaning that low-quality males do engage in contests but also pursue other tactics, adaptive capacity is reduced and evolutionary rescue, where a population is able to adapt to a changing environment, becomes less likely. This is because the use of the ART allows low-quality males to contribute more maladaptive genes to the population than would happen otherwise. When the ART is fixed, however, such that low-quality males will only use the alternative tactic and do not engage in contests, we find the opposite: adaptation happens more quickly and evolutionary rescue when the environment changes is more likely. This surprising effect results from an increase in the mating success of the highest quality males who face many fewer competitors in this scenario—counterintuitively, the presence of males pursuing the ART increases reproductive skew towards those males in the best condition.

## Introduction

Organisms usually exist in habitats that they are reasonably well adapted to. When the environment changes such that those organisms suffer substantial reductions in fitness there are several possible outcomes: the population in question might become extinct, or the organisms might migrate to or otherwise colonise new environments which have become more suitable to them. A third possibility is ‘evolutionary rescue’: the phenomenon whereby the affected population evolves rapidly enough to become adapted to the new environment and thus is able to persist despite the environmental changes (Bell, 2017). Evolutionary rescue will only occur when the rate of adaptation is sufficient to offset the fitness reductions arising from environmental change. Given that the environment is changing rapidly in almost every habitat on the planet and many populations likely have limited ability to migrate due to geographical barriers or habitat fragmentation, understanding the factors that determine how fast adaptation can occur is necessary to allow prediction or management of populations’ responses to changing environments.

Over recent decades it has become clear that mating systems can be important determinants of adaptive capacity for animal populations—there is now firm evidence from laboratory studies that strong sexual selection can enhance adaptation and therefore the probability of evolutionary rescue (Cally et al., 2019; Godwin et al., 2020; Jarzebowska and Radwan, 2010; see Parrett et al., 2019 for a field study). These positive effects of strong sexual selection are a consequence of sexual signals expressed by (usually) males being condition dependent (Dougherty, 2021; Rowe and Houle, 1996; Tomkins et al., 2004), meaning that they respond disproportionately to the overall health and well-being of the bearer. If females base their mating decisions on these signals, or if male-male contests are determined on the basis of their expression, then males that are well-adapted to the new environment, and therefore in better condition, will gain a high proportion of matings and father a higher proportion of the offspring in the next generation than would otherwise occur, leading to more rapid adaptation and potentially a greater probability of evolutionary rescue (Agrawal and Whitlock, 2012; Lorch et al., 2003; Martínez-Ruiz and Knell, 2017; Whitlock, 2000).

Mating systems are, of course, greatly diverse and strong sexual selection with either female choice or intrasexual contests leading to high reproductive skew in favour of a few high-quality males is only one of many possible variants, each of which could have differing effects on the rate of adaptation and the potential for evolutionary rescue. Here we consider the potential role of one widespread variant, whereby males adopt alternative reproductive tactics (ARTs), either competing with rivals for access to mates (variously called ‘major’, ‘fighter’, ‘bourgeois’ or ‘guarder’ males) or attempting to acquire matings by means other than direct competition (‘minor’, ‘scrambler’, ‘parasitic’ or ‘sneak’ males) (Engqvist and Taborsky, 2016; Gross, 1996; Neff and Svensson, 2013; Oliveira et al., 2008). ARTs are diverse and distributed across the animal kingdom. Some, such as those found in the ruff *Philomachus pugnax* (Lamichhaney et al., 2016; Lank et al., 1995) and the side-blotched lizard *Uta stansburiana* (Sinervo and Lively, 1996) are largely determined by genetic polymorphisms associated with specific mating phenotypes, with the polymorphism being maintained by frequency-dependent selection. Others, however, are polyphenisms, whereby any single genotype can develop into multiple phenotypes depending on their environment. In the dung beetle *Onthophagus taurus*, for example, males either develop into horned ‘major’ males or effectively hornless ‘minor’ males depending on food availability when larvae (Hunt and Simmons, 1997; Moczek and Emlen, 1999). The horned major males guard females in tunnels and will fight other major males who intrude, but minor males do not fight and instead attempt to acquire matings using non-aggressive tactics including mating with females when they leave the burrow to collect dung or passing a guarding male without fighting to access the female beetle (Moczek and Emlen, 2000). In some cases morph determination is a complex product of both genetic and environmental cues: in the case of the bulb mite *Rhizoglyphus robini* males develop into either ‘fighter’ morphs, armed with an enlarged and sharp third pair of legs used to kill other males in contests, or ‘scrambler’ morphs which avoid contests and seek matings by other means. In this species development into a specific morph is partly heritable (Radwan, 2003, 1995), shaped by genome-wide variation (Parrett et al., 2022) and with considerable additive genetic variance (Parrett et al., 2023), but is also strongly influenced by size in the final instar, which is itself determined by both maternal investment in the egg and by food availability (Smallegange, 2011).

In general, the polyphenic expression of morphs which engage in alternative tactics is thought to be an example of conditional expression, whereby individuals which are small, weak or otherwise of low status adopt alternative, non-aggressive mating tactics because these tactics lead to higher fitness for them. Large, strong or otherwise high-status males, on the other hand, gain the greatest fitness benefits from competing aggressively (Gross, 1996; Hunt and Simmons, 2001; Tomkins and Hazel, 2007). Since the generally accepted explanation for the mechanism by which sexual selection enhances adaptation is based on the idea that males in good condition (high status) acquire the majority of matings, it might be expected that this effect should be disrupted by the expression of ARTs since these increase the mating success of males in poor condition, weakening selection against deleterious alleles or for beneficial ones.

Here, we develop and analyse an individual-based model (IBM) of populations evolving in a changing environment, with males able to adopt either a competitive or a non-competitive strategy on the basis of their condition—thus we consider only the status dependent case of ARTs and not the genetically determined frequency-dependent option which is very different. IBMs are a useful approach to explore complex evolutionary processes which interact with demography because they allow individual variation to be explicitly included (DeAngelis and Mooij, 2005). This IBM is based on an earlier model (Martínez-Ruiz and Knell, 2017) which simulated a population evolving under variable environments with different degrees of sexual selection and condition dependence. By adapting the model to include alternative mating tactics it is possible to investigate whether the probability of evolutionary rescue is altered by the use of these tactics by males under a variety of scenarios with different values for, for example, the overall size of the population, the average fecundity and the probability of low-status males adopting the alternative tactic.

We distinguish between ‘simultaneous’ and ‘fixed’ ARTs (Taborsky et al., 2008) where simultaneous ARTs can be expressed at the same time as the ‘normal’ tactic by the same individual, as in three-spine sticklebacks where nest holding males will also engage in sneak tactics, (Rico et al., 1992) and fixed ARTs are those where one individual is only able to express one tactic, such as the *O. taurus* and *R. robini* examples given above. Population persistence under directional environmental change and the speed of return to a near-optimal level of adaptation after a step-change in the environment are both modelled.

## Results

### Directional change

In our model the environment is modelled as a single continuous variable and an individual’s optimal environmental value is determined by a single continuously varying variable called *environmental genotype*. An individual’s *condition* is a value between 0 and 1 determined by the difference between *environment* and *environmental genotype*, and whether an individual pursues an ART is determined by whether *condition* is below a *threshold* value. If the *threshold* is 0 then no males will follow the ART and if it is 1 then all will.

Figure 1 shows a typical output from the model when the environment changes in a directional fashion and alternative tactics are simultaneous, meaning that males adopting these tactics participate in contests for females but also try to acquire matings by other means. While the environment remains roughly constant the population fluctuates around the carrying capacity. There is some skew in the adult sex ratio as a consequence of the extra mortality cost experienced by major males and the proportion of minor males in the population is roughly a third. At *time =* 125 the environment starts to undergo directional change. The population does evolve in response, as can be seen by the increasing value of *environmental genotype* (the variable that determines the optimal value for the environment for each individual) in panel B but the rate at which the population can adapt is slower than the rate of change of the environment and the median condition of the population declines (panel C), leading to both an increased death rate and lower reproduction by females. Consequently the population declines until extinction occurs after roughly 400 time steps. As the median condition of the population declines the proportion of minor males increases which will further reduce the adaptive capacity of the population since more females will be mating with these poor-condition males. Finally, note that the sex-ratio skew in the adult population is reduced when the proportion of minor males is high. These males do not express a sexual signal and therefore do not experience an increased probability of mortality.

**Figure 1.**
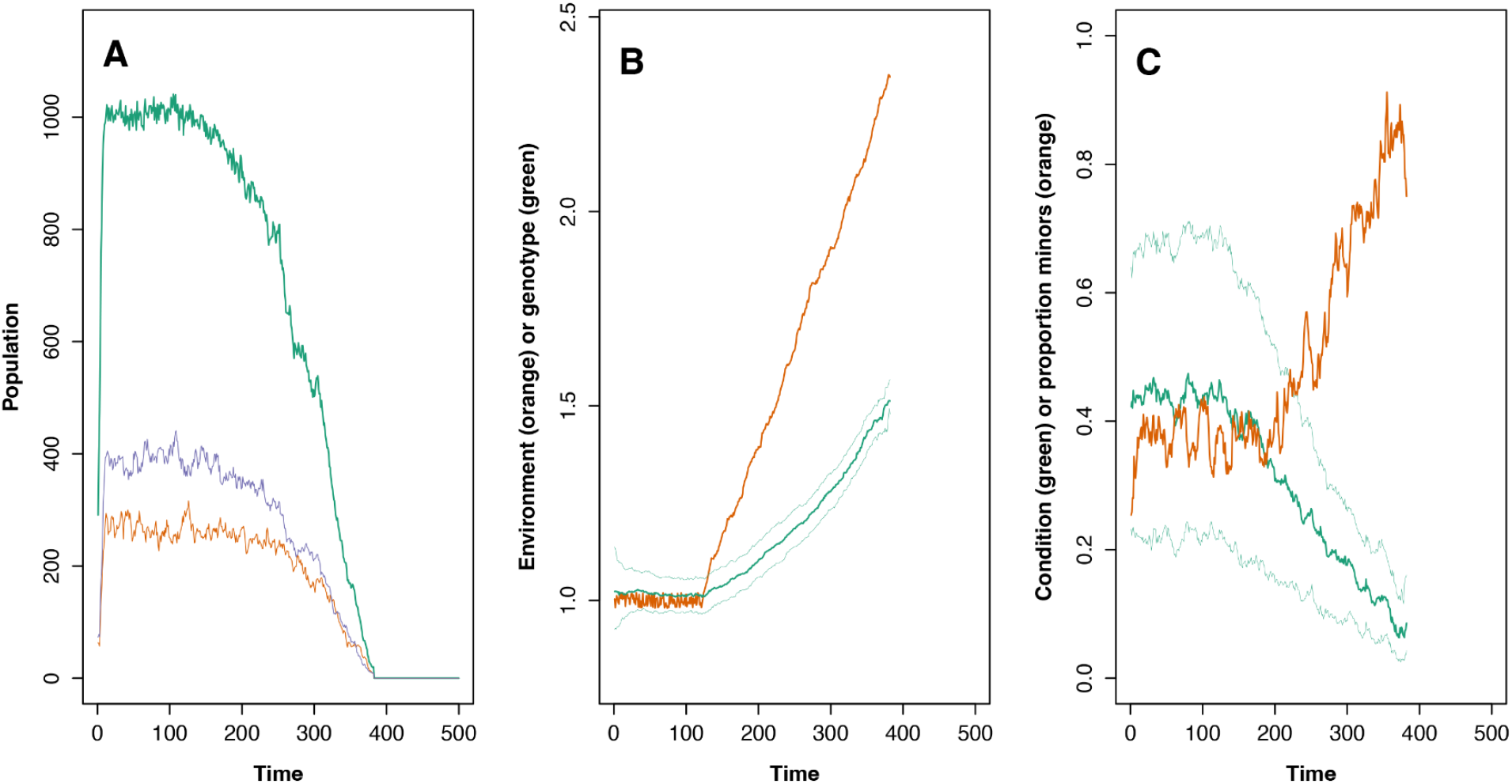
Model output showing extinction when male alternative tactics are simultaneous. A: Total population (green) plus the populations of mature females (purple) and males (orange). B: The value of the *environment* variable at each time step (orange) plus the median and quartiles of the *environmental genotype* variable (green). C: The median and quartiles of *condition* plus the proportion of males adopting the minor strategy (orange). Parameter values for the simulation are *K = 1000, threshold = 0*.*25, max_offspring = 6, group_size = 6, ART_success* = 0.5 and *β = 2*.

Figure 2 shows the proportion of simulations where extinction took place for a variety of parameter values when the rate of directional change was set to a single value which was sufficient to cause extinction in all simulations when mating was random—in this model this occurs when either the mating group size or the threshold condition value (the value for *condition* which determines weather males follow the ART or not) for pursuing the alternative mating tactic threshold is 1. If there is relatively strong sexual selection and no males pursuing alternative strategies (mating tactic threshold = 0, mating group size is >1 and *β* >1 then evolutionary rescue can occur, especially when the population is large and sexual selection is strong (mating group size is 10 and *β* = 2 or 4), in which case all populations are able to persist over the course of the simulation. The inclusion of some males pursuing alternative tactics (mating tactic threshold = 0.25 or 0.5), however, negates this effect of sexual selection and the probability of evolutionary rescue of these populations is reduced substantially.

**Figure 2:**
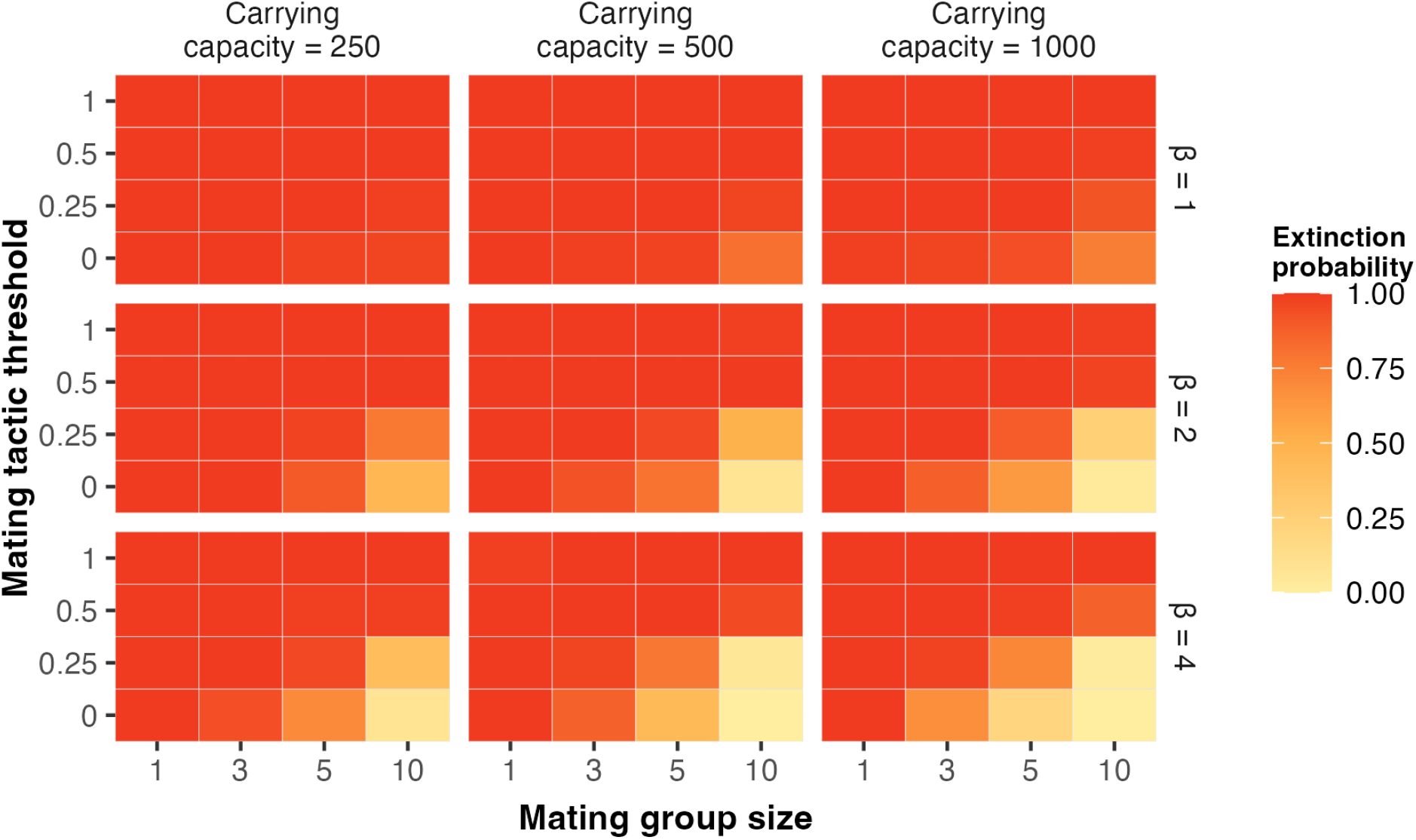
heatmap showing the probability of extinction under directional environmental change when alternative tactics are simultaneous, calculated from 100 simulations running for 500 time steps for each combination of parameter values. The rate of directional change was set to 0.005 for all simulations. Other parameter values: *max_offspring* = 6, *ART_success* = 0.5.

By contrast with the situation for simultaneous alternative tactics, when the alternative tactics were fixed and minor males did not compete in groups for access to females, the presence of males pursuing alternative tactics increased the probability of evolutionary rescue. Figure 3 shows a set of outputs from the model when alternative tactics are fixed and the population persists. In this case the median value of *environmental genotype* is able to keep pace with the increasing value of *environment* and the population persists, albeit at a size far below the carrying capacity and with most individuals in poor condition.

**Figure 3.**
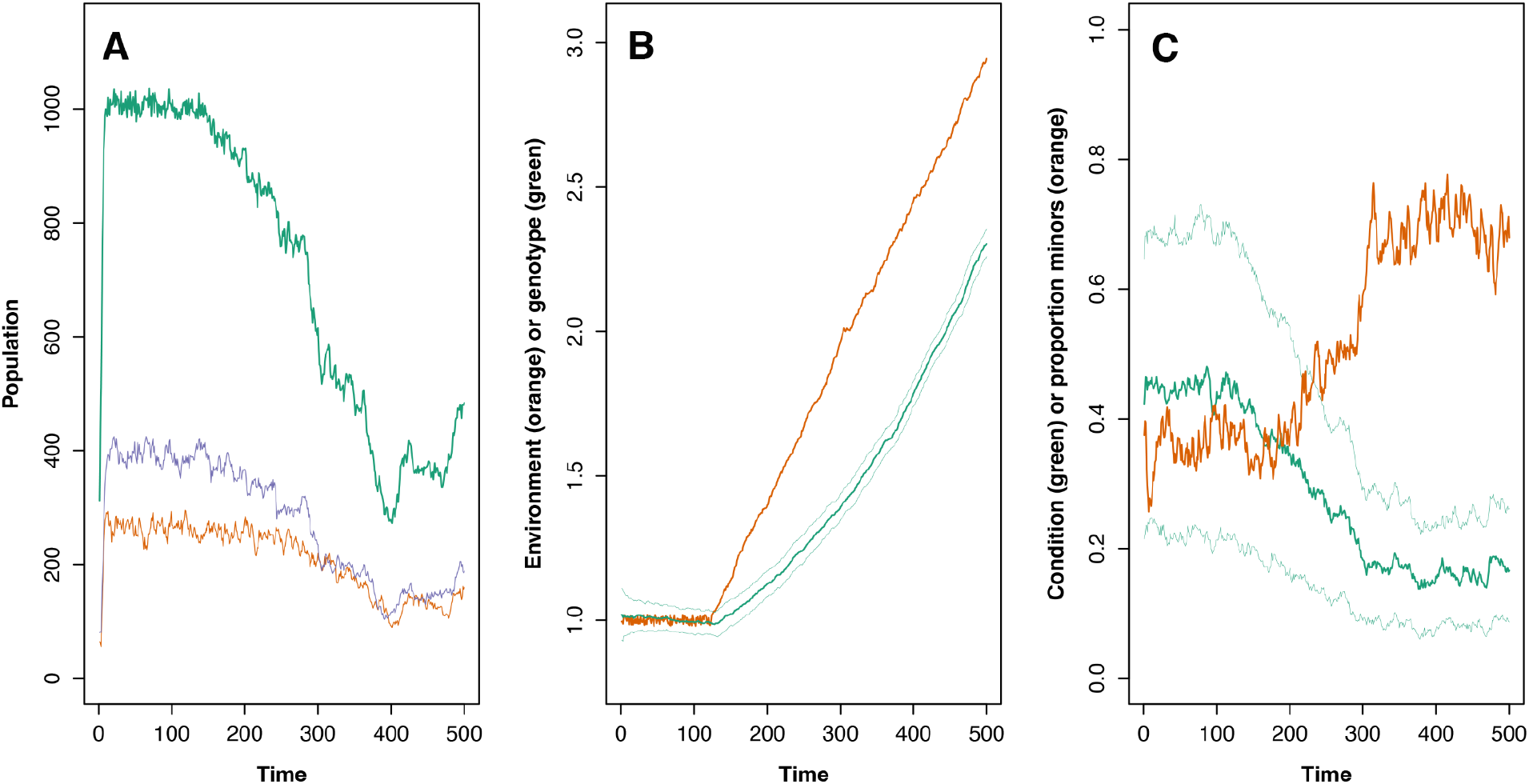
Model output showing extinction when male alternative tactics are simultaneous. A: Total population (green) plus the populations of mature females (purple) and males (orange). B: The value of the *environment* variable at each time step (orange) plus the median and quartiles of the *environmental genotype* variable (green). C: The median and quartiles of *condition* plus the proportion of males adopting the minor strategy (orange). Parameter values for the simulation are *K =* 1000, *threshold =* 0.25, *max_offspring =* 6, *group*.*size =* 6, *ART_success =* 0.5 and *β = 2*.

Figure 4 shows the proportion of simulations where the population became extinct for a range of parameter values with fixed alternative tactics. As with figure 2, when the mating group size or the condition threshold for pursuing alternative tactics is 1 then mating is random and all populations become extinct, and when there are no males pursuing the alternative tactic (mating tactic threshold = 0) then sexual selection can lead to evolutionary rescue, especially when sexual selection is strong (i.e. larger values of *β* and mating group size) and the population is larger. When some lower condition males pursue the alternative tactic and do not compete with the other males, however, evolutionary rescue is more likely, occurring at lower values of *β* and mating group size than when males do not pursue the alternative tactics, and in smaller populations.

**Figure 4:**
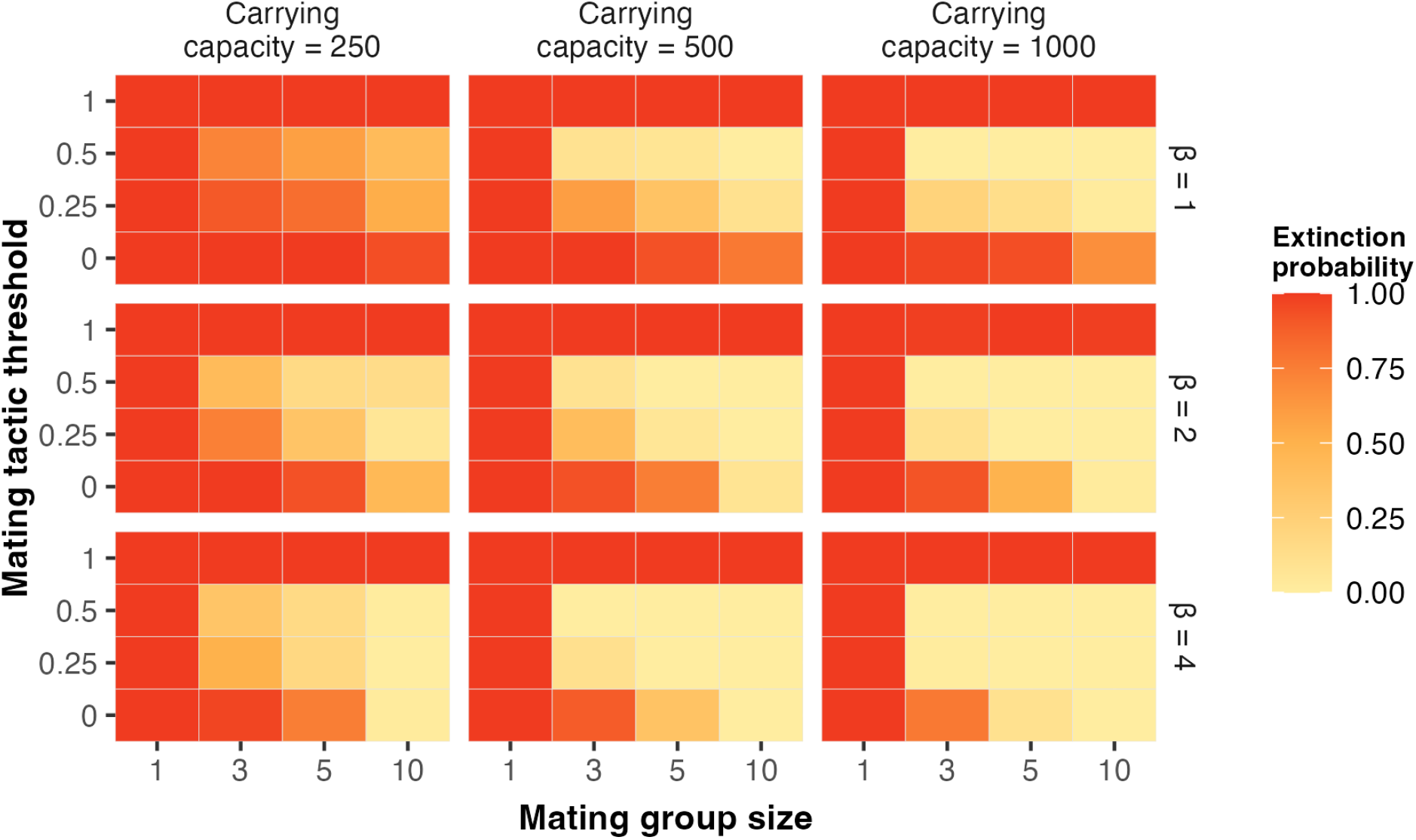
heatmap showing the probability of extinction under directional environmental change when alternative tactics are fixed and males pursuing those tactics do not compete with other males in groups, calculated from 100 simulations running for 500 time steps for each combination of parameter values. The rate of directional change was set to 0.005 for all simulations. Other parameter values: *max*.*offspring* = 6, *ART_success =* 0.6, *s* = 0.5.

### Environmental step-change

Median return times for simulations that underwent an environmental step change are shown in figure 5, with the return time here being the number of time steps after the step change before the upper quartile of the distribution of values for the *environmental genotype* became equal to or greater than the value for *environment*. The strength of sexual selection increases with both the mating group size and the parameter determining the strength of female choice or the advantage of higher quality males in contests within a group (*β*) and the return time declines as both of these increase, indicating faster adaptation. When the alternative tactics are simultaneous the presence of males following alternative tactics leads to slower return times and when males are more likely to follow the ART (*threshold* = 0.5 rather than 0.25) there is a greater increase in return time and therefore slower adaptation. When the ART is fixed, however, the inverse is found and the presence of males following the ART leads to shorter return times and therefore faster adaptation. Thus, consistent with the results from the directional change case above, simultaneous ARTs reduce adaptation whereas fixed ARTs increase it.

**Figure 5:**
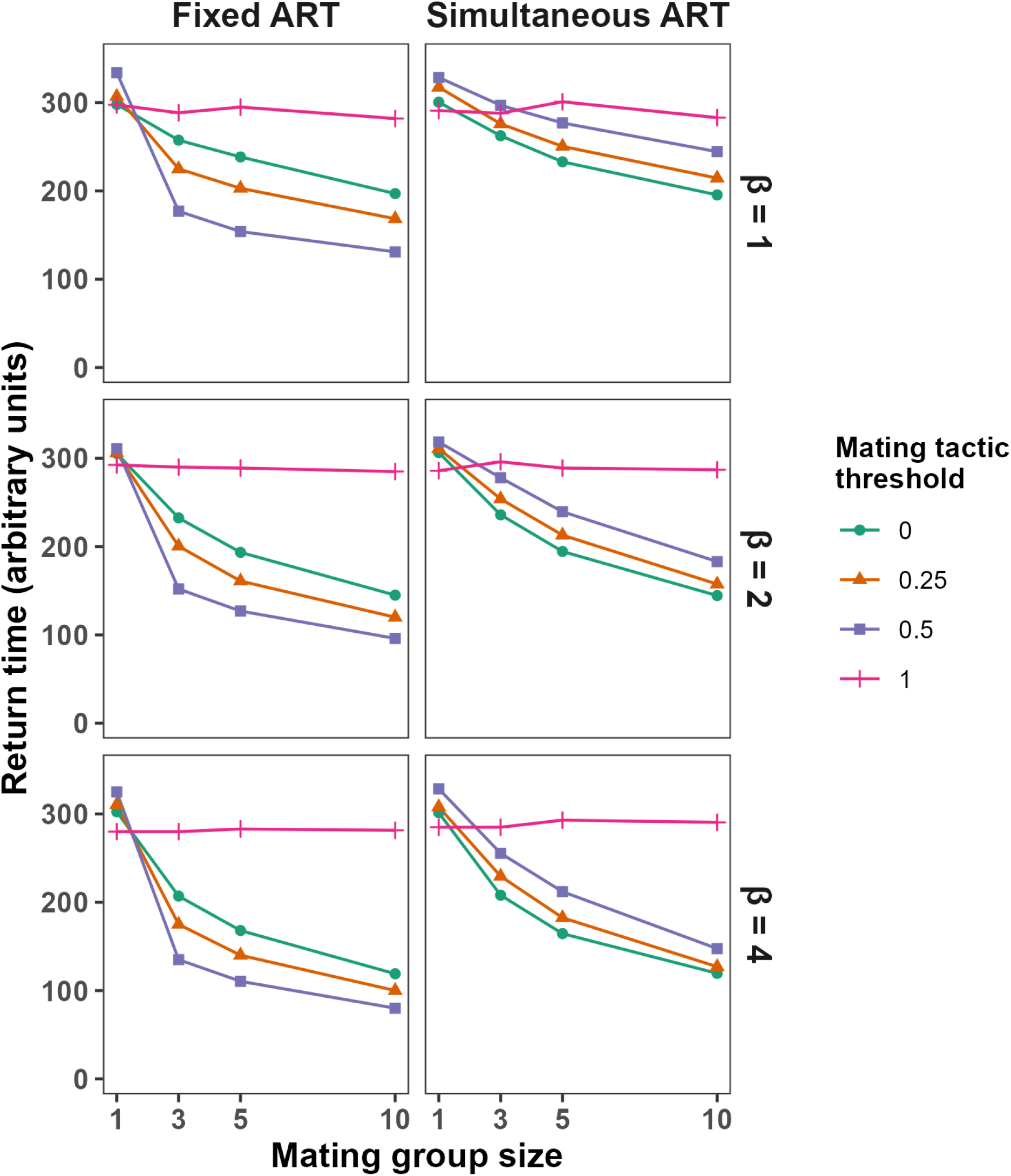
median return times for populations following an environmental step change. Return times are the number of time units before the upper quartile of the distribution of *environmental genotype* is equal to the value of *environment*. When the mating tactic threshold = 1 all males follow the ART meaning that there is random mating in the simulation and males do not express sexual signal traits. Each value is the median from 100 simulations, and the carrying capacity was set at 500 for all simulations.

When the mating tactic threshold = 1 then all males follow the ART and so the return time is unaffected by the strength of sexual selection. Interestingly when sexual selection is very weak or non-existent (group size = 1) then the return times for the cases when all males follow the ART are actually lower than the alternatives. This is most probably because in this model males following the ART do not express the sexual signal and so do not pay a cost for doing so, whereas even when the group size = 1 males not following the ART will pay the cost for expressing the signal. Well-adapted, high quality males will therefore have a somewhat longer life in the simulations where all males follow the ART and will consequently produce more offspring.

### Reproductive skew with simultaneous and fixed ARTs

Why do simulations with fixed ARTs show faster adaptation and an enhanced probability of evolutionary rescue? In this case the low-condition males are excluded from the mating groups where males compete for matings. If all males are allocated to these groups, as when there are no males pursuing alternative tactics, or when the alternative tactics are simultaneous, then purely by chance some groups will have no high-condition males, giving the low-condition males in those groups an opportunity to mate. If low-condition males do not compete then all the groups of competing males will consist of high-condition males, enhancing the probability of evolutionary rescue because under these conditions all the females will mate with a relatively high-condition male, who most likely will have a genotype that is well matched to the changing environment.

To further explore this we calculated the amount of reproductive skew in males for each timestep for simulations with simultaneous and fixed males. Reproductive skew was calculated as the degree of skewness in the distribution of offspring per male using the moments package in R (Komsta and Novomestky, 2015). These data are shown in figure 6. While the amount of reproductive skew is weakly negatively correlated with the proportion of minor males in the simultaneous case, when alternative tactics are fixed there is a clear positive relationship between reproductive skew and the proportion of males pursuing the alternative tactic.

**Figure 6:**
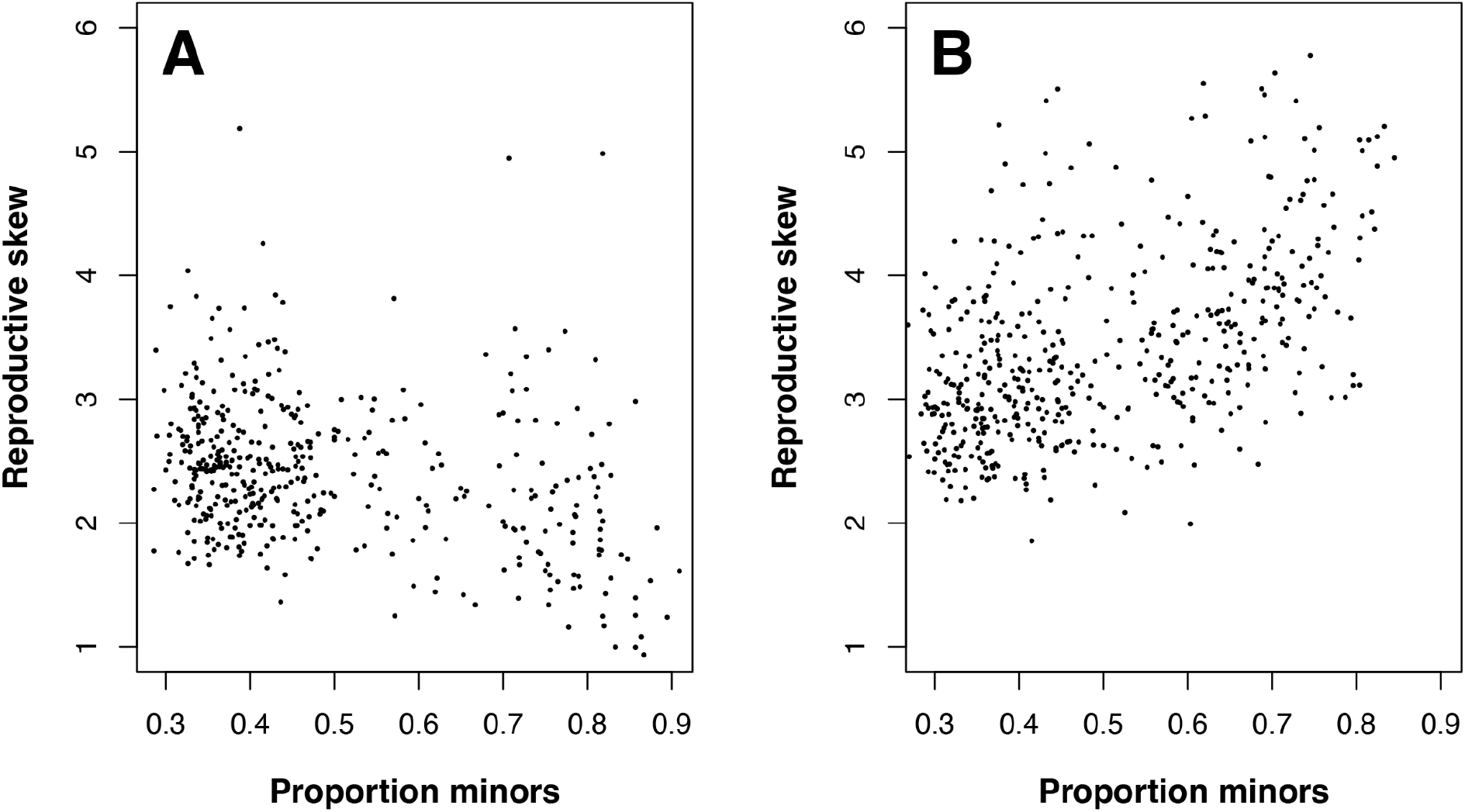
reproductive skew in males plotted against the proportion of minor males in the population for each timestep of a simulation running for 500 timesteps. A: alternative tactics are simultaneous and male reproductive skew decreases slightly with the proportion of minor males. B: alternative tactics are fixed, male reproductive skew increases with the proportion of males following the minor strategy. Other parameter values: *K* = 500, *ART_success* = 0.5, *max_offspring* = 6, *group_size =* 6, *threshold =* 0.25 and *β* = 2.

## Discussion

Alternative reproductive tactics (ARTs) are found across the animal kingdom, so if they have an impact on adaptation and evolutionary rescue it is important that we understand this. On the basis of this model we find that whether the ART is simultaneous or fixed is a crucial determinant of the population-wide evolutionary impact of the ART. When ARTs are simultaneous the capacity of the population to adapt is reduced and evolutionary rescue is less likely when the environment is changing. When ARTs are fixed, however, and males pursuing the alternative tactic do not engage in contests with other males, the adaptive capacity of the population is enhanced and evolutionary rescue becomes more likely.

The reduced adaptive capacity of populations when the ART is simultaneous is intuitively understandable. Poorly adapted males, with a phenotypic optimum which is far from the existing value, will be in poor condition and unlikely to win contests with better adapted males who are in better condition. By adopting the ART, these males will acquire matings that would otherwise be unavailable to them, and will father offspring with maladapted alleles who would otherwise most likely be fathered by better adapted males, inhibiting the process of adaptation at the population scale. These negative effects will likely be greatest in systems where females are unable to exert any choice over mating rates with males adopting the ART. For example, male guppies (*Poecilia reticulata)* perform courtship displays in order to solicit matings with females. If, however, females are unreceptive then males may adopt a coercive mating strategy in which females have little choice over and these coercive copulations have been shown to transfer a considerable amount of sperm, possibly undermining female choice in this system (Pilastro and Bisazza, 1999). If the coercive ART imposes a cost to females via traits that increase male fitness at the expense of females, then again we would anticipate increased population level negative effects when males adopt the ART, in a similar way to when harmful male traits are themselves condition-dependent (Flintham et al., 2023). Thinking more widely, increasing polyandry generally might reduce adaptive capacity in a similar fashion and this might explain the pattern found in fossil ostracods by (Martins et al., 2018) where increasing sexual dimorphism arising from larger sperm pumps, presumably being associated with increased sperm competition, was found to be closely correlated with increased extinction risk.

The enhanced adaptive capacity of populations when the ART is fixed, on the other hand, does not seem intuitively obvious. As discussed in the results, this effect occurs because once the males following the ART are removed from competition, the average condition (and therefore degree of adaptation) of males undergoing competition will be higher, leading to greater reproductive skew towards the males in the best condition and consequently a greater representation of alleles conferring better adaptation in the next generation. This effect is robust and persists even when males pursuing ARTs have enhanced probabilities of acquiring matings and fathering offspring (see supplementary material).

This surprising result for fixed ARTs is, of course, a theoretical outcome from a simulation model, and obviously it is necessary to ask whether we should expect this mechanism to operate in real animal populations. There are no empirical studies that we are aware of that would provide a direct hypothesis test, but we can at least examine the assumptions of the model and consider whether there is any further evidence that might help us to address this. The strength of this effect relies on two assumptions in the model: firstly, that the ART is fixed and secondly that females will continue to mate with males which are not following the ART even when such males are rare. The first assumption, that following the ART is fixed and that males pursuing this tactic do not engage in contests is certainly valid for many systems where ARTs are used—minor dung beetles do not invest in the horns that are necessary for contests with majors (Simmons et al., 2007), for example, and scrambler morphs of the mites *Sancassiana berlesei* and *Rhizoglyphus robini* suffer high mortality from fights (Radwan and Klimas, 2001) and therefore avoid fights with fighter morphs. The second assumption, that even when good-condition males which are not following the ART are rare, all the females in the population will continue to mate with these males in the same way as when they are common, is weaker. If we consider a female-choice system then when such males are very rare some females are likely to mate with low-quality males if they are unable easily to find appropriate high-quality males. There is also evidence that the strength of female choice itself can be altered by both prior experience (Bailey and Zuk, 2008; Macario et al., 2017) and the female’s condition (Hunt et al., 2005), so choice overall might become weaker in a stressed population because the females are in poor condition themselves and because if the female has little or no prior experience of high-quality males she is likely to be less choosy than otherwise. Similar arguments can be made about systems where the males engage in contests to monopolise females. In the extreme then, and in populations that are especially stressed by the changing environment the strength of the effect we have found is likely to be reduced, but it will not be eliminated altogether. In species where males compete to guard rare resources that are essential or desirable for female reproduction, such as oviposition sites, then even when only a few high-quality males are present these will still enjoy a higher mating success than otherwise because of the lack of rivals.

Does the phylogenetic distribution of ARTs tell us anything about whether they might enhance or reduce persistence of a species? If the presence of males following ARTs reduces species persistence we might expect to see a ‘twiggy’ distribution of ARTs, with species expressing ARTs being found at individually or in small clades at the tips of phylogenies, but with no large taxa expressing them, as has been suggested to be the case for asexual reproduction (Maynard Smith, 1978; Schwander and Crespi, 2009). An analysis of the phylogenetic distribution of ARTs across the animal kingdom is not possible with current knowledge but we can note that within the Coleoptera there are a number of large clades with males that seem mostly to pursue ARTs. McCullough et al., (2015) examined the static allometry of the horns carried by males of 31 species from the Dynastinae (Hercules beetles) and found discontinuous relationships suggestive of ART expression in 30 of them. Within the Lucanidae (stag beetles), Matsumoto and Knell (2017) found evidence for complex polymorphisms, including species with three and even four male morphs, in all six species of *Odontolabis* examined. Within the Scarabaeinae (true dung beetles) the picture is somewhat more complex because, in the genus *Onthophagus* at least, male weaponry is known to be evolutionarily labile such that horns are rapidly gained and lost over evolutionary time (Emlen et al., 2005). Nonetheless, when the horned dung beetles are examined the great majority of species seem to exhibit male polymorphism that is most probably associated with males following either a ‘guard’ or a ‘sneak’ strategy (Parrett et al., n.d.; Simmons et al., 2007). The existence of clades of animals where most or all males seem to pursue ARTs suggests that in the beetles at least the use of these tactics is not a significant contributor to the probability of extinction and is possibly enhancing species persistence. The behaviour of the minor males has only been studied in detail in a relatively small number of species from the Scarabaeinae but it is notable here that these minors all seem to follow something approximating to a fixed strategy—in *Onthophagus acuminatus* and *O. taurus*, for example, minor males will enter burrows and remain there with females but if challenged by a major male are unable to compete and resort to ‘sneak’ behaviour to acquire matings (Emlen, 1997; Moczek and Emlen, 2000). Future work might focus on ART distributions in fish: both fixed and simultaneous ARTs are found within the ray-finned fish (Taborsky, 2008, 1998) and have been shown to have multiple independent origins (Mank and Avise, 2006). Addressing the distribution of each type of ART across this group may provide a direct test of the possibility that ART type influences extinction risk

As a final point, we should point out that this model is only appropriate for considering the impact of some types of ART on the adaptive potential of a population. Alternative reproductive tactics are diverse and we have not considered, for example, sequential ARTs where the reproductive tactic changes through the life of an animal. These are common in fish and mammals (Oliveira et al., 2008) and might have different effects on adaptive potential. ARTs that are based on genetic polymorphisms are very different from the polyphenic type modelled here, and whether these might alter adaptive potential is currently an open question.

## Methods

The model is coded in R (R Development Core Team, 2022) and is an adaptation of the one described in (Martínez-Ruiz and Knell, 2017). It simulates the dynamics and evolution of a spatially homogeneous, age-structured population of animals. The model proceeds as a series of discrete time steps of arbitrary length. All simulations described here were run for 500 time steps or until the population became extinct, whichever was shorter. Multiple simulations were run in parallel using the doparallel (Microsoft Corporation and Weston, 2022a) and foreach (Microsoft Corporation and Weston, 2022b) packages running in R version 4.3.0 on a 2021 MacBook Pro with a 10-core M1 Pro CPU and 32GB RAM.

The environment is described by a single continuous variable *environment* which is best thought of as representing an environmental variable such as temperature or pH. The value for *environment* is initially set at 1 and two different treatments were used: directional change and step-change. For directional change, the environment was held constant aside from a small amount of random variation for the first 25% of the time steps. Following this and for the rest of the simulation the environment changes in a directional way, with a value drawn from a normal distribution with *μ* equal to a parameter *directional rate* and *σ=0*.*005* added to the value at each time step. For the step-change treatment the environment was allowed to remain constant aside from a small amount of random variation, but at time point 100 the value of *environment* had 0.33 added. This change is not sufficient to make the population become extinct, but populations exposed to such an environmental change will be some distance from being optimally adapted once the change has happened.

Populations are started with 200 individuals which are allocated the following characteristics: *sex* (male or female, allocated at random); *age* (a value from 1 to 10, drawn at random from a uniform distribution); *environmental genotype*, which determines how well that individual is adapted to the environment (drawn for each individual from a normal distribution with *σ = 0*.*25* and *μ =* the starting value for the ‘environment’ variable) and *resource*, which describes how each individual is able to acquire resources independently of environmental effects (drawn at random from a uniform distribution with minimum 0 and maximum 1). Including *resource* in the model here adds an element of environmental variance to the expression of the sexual trait which in real systems will be determined by both environmental and genetic variance (Singh and Agrawal, 2022).

Each individuals condition is then calculated as:

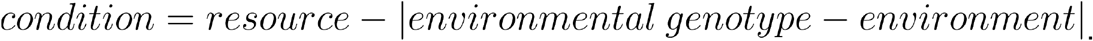

Individuals with *condition <0* die, so *condition* is a value between 1 and zero for all living individuals.

The variable *threshold* determines whether an male with a given value for *condition* will develop into a major male (aggressive strategy) or a minor male (non-aggressive strategy). The strategy a male follows is determined at *age* = 1 and then fixed for the lifetime of that individual, so this aspect of the model is representative of alternative reproductive tactics in examples such as bulb mites and dung beetles and salmon where the alternative tactic is determined before sexual maturity. If *threshold =* 0 all males follow the aggressive strategy. Major males over the age of maturity (set at 2) express a display trait (*display*) which is equal to their *condition*. Minor males do not express such a trait. Two versions of the ART were used, one where the strategy is fixed (*sensu* Taborsky et al., 2008)), and minor males do not attempt to compete for matings, and one where the strategy is simultaneous and minor males will compete for matings but will also try to acquire matings by other means.

Every time step, the *age* for each individual is incremented by 1. Following this, the probability of dying is calculated for each individual as:

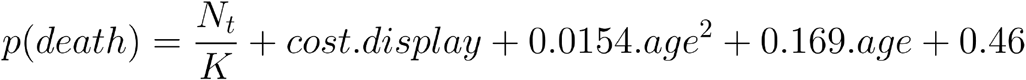

Where *N*_*t*_ *= population size, K* determines the maximum population size, *cost* is the cost of bearing the display trait (note the display trait is set to zero for females, minor males and males below the age of maturity) and the quadratic equation gives higher mortality for young and old individuals.

For mate allocation, males are divided into groups with size determined by a parameter *g*. If the ART is fixed then only major males are allocated to these groups, if simultaneous then all males above the age of maturity are allocated to mating groups. The males in each group are ranked on the basis of the value of their display trait, and each female of reproductive age is allocated to one of the groups of males.

Each female will mate with one male from the group she is allocated to. The probability of mating is calculated for each male in a group on the basis of his ranking and the value of a parameter *β* as follows:

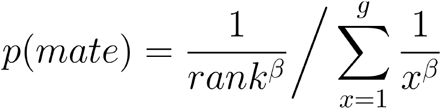

*β* controls the relationship between display trait expression and mating—this can be thought of as either the strength of female preference or the strength of the advantage in contests between males given by the display trait, with higher values of *β* meaning that males with large display traits have a greater advantage.

Following the allocation of mates via contests between males, each female can also mate with a randomly selected minor male with a probability equal to the proportion of minor males multiplied by a parameter *ART_success*, which allows the success of the minor strategists to be adjusted. Each female then produces a number of offspring equal to the maximum offspring multiplied by her condition, rounded to the nearest integer. For females who mate with both a major and a minor male, paternity for each offspring is allocated at random with equal probability for both males.

The *environmental genotype* is modelled as a quantitative, polygenic trait controlled by many alleles so each offspring has an *environmental genotype* equal to the mean of its parents’ *environmental genotype* value plus a value drawn from a normal distribution with *mean = 0* and *σ* equal to a parameter called *mutation*.

For simulations run with directional change the response variable used was whether the population became extinct over the course of the simulation. When the environmental change was a step-change the response variable used was the number of time units after the change before the upper quartile of the distribution of values of *environmental genotype* first became equal to or greater than the *environment* variable. This gives an indication of how long it takes the population to adapt to the new value of *environment*.

The supplementary material contains details of a further set of analyses exploring the effect of variables including the ART type, mating group size, *β* and *ART_success* in determining the Critical Rate of Environmental Change (Chevin et al., 2010) for these simulated populations. The model code and code to replicate the simulations and analyses is [archived on GitHub](https://github.com/rjknell/Alternative_tactics_model).

## Supporting information

Supplementary results

