## Supplementary results for "Alternative reproductive tactics and evolutionary rescue"

### Alternative reproductive tactics and evolutionary rescue: supplementary material

#### Critical rates of environmental change and the strength of sexual selection

To further explore the way that the presence of males using alternative tactics affects evolutionary rescue under directional environmental change, we calculated the Critical Rate of Environmental Change (Chevin et al., 2010)—the rate of environmental change above which a population cannot adapt and persist—for a variety of parameter combinations . Because of the stochasticity in these models there is not a single rate above which a population will persist, so the critical rate was approximated as the rate of environmental change at which 50% of simulations became extinct. This was estimated by running 100 simulations at a range of values for the rate of change (initially 0.002 to 0.008 in steps of 0.001 and then further values from 0.00325 to 0.00575 in steps of 0.0025 in order to achieve more precise estimates) and fitting a generalised linear model with binomial errors to the data on extinction with rate of change as an explanatory variable. The rate of change at which 50% of populations were predicted to become extinct was then extracted from each model using the *dose.p()* function of the MASS package running in R (Venables and Ripley, 2002).

Figure S1 shows the effect of including fixed or simultaneous alternative tactics and the size of the mating groups for three values of $\beta$, the parameter that controls the probability of the highest ranked male acquiring a mating in a particular mating group. Both larger group sizes and higher values of $\beta$ will lead to stronger sexual selection in the population, and values for the critical rate are highest when both of these values are high. For simultaneous alternative tactics, which, as we have seen, tend to decrease persistence in changing environments the effect size (the magnitude of the decrease in the critical rate) is largest when sexual selection overall is strong, with the largest decreases in the critical rate of environmental change when *β* = 4 and *group.size* = 10. There is also an interaction between the strength of sexual selection and *threshold*, the parameter that specifies the value of *condition* below which a male develops into a minor male, such that the critical rate of environmental change decreases much more with strong sexual selection when *threshold* = 0.5 than when *threshold* = 0.25.

When the alternative tactics are fixed then the effect of introducing alternative tactics is positive, meaning that the critical rate of environmental change is greater when males in the population pursue alternative tactics. When the threshold for pursuing alternative tactics (*threshold*) is set at a *condition* of 0.25 then introducing the fixed ART leads to a consistent increase in the critical rate of roughly 0.001 no matter what the strength of sexual selection, but when *threshold* is 0.5 the extra increase in the critical rate arising from the alternative tactic is reduced when sexual selection is strongest, in other words when *group_size* is 8 or 10 and *β* is 4. The reason for this is not clear but we suggest that ultimately adaptation in the population will be limited limited by other factors such as the population size and the average magnitude of mutations: if the environment is changing very rapidly then there will simply not be variation being added to the *environmental genotype* to allow adaptation no matter what the mating system.

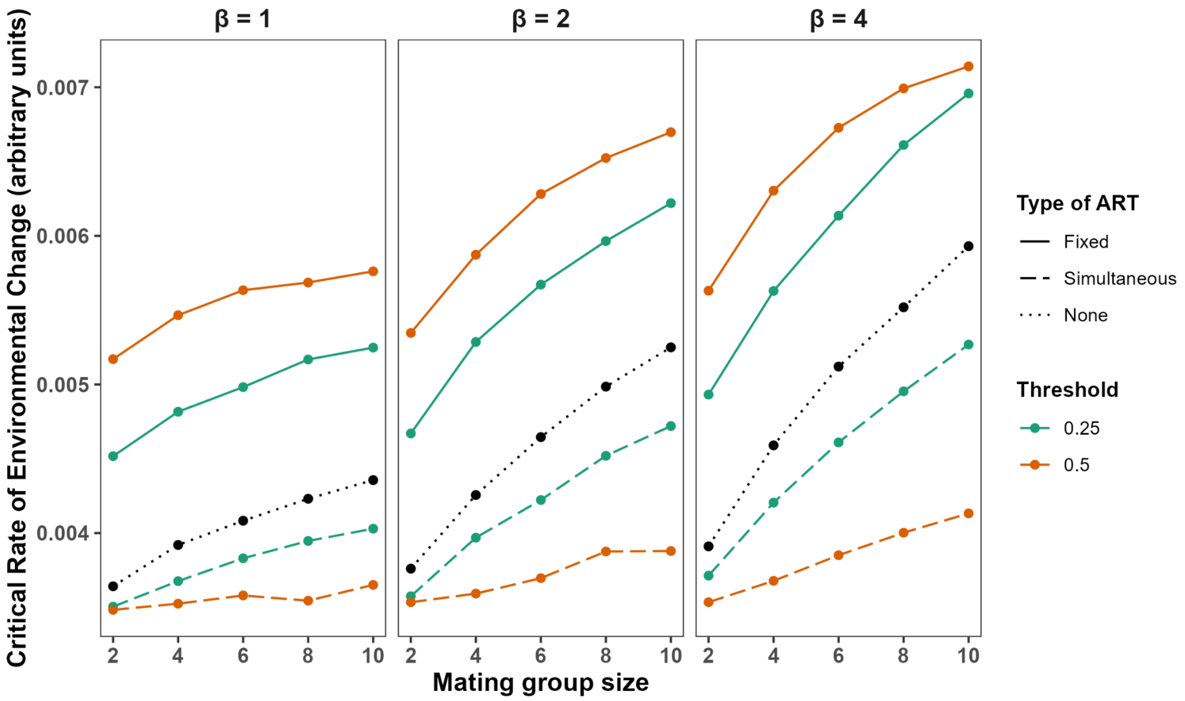


**Figure S1:** variation in the critical rates of environmental change with increasing strength of sexual selection for populations without ARTs and for populations with males expressing fixed or simultaneous ARTs. The two parameters that between them determine the strength of sexual selection in a simulated population are *group_size*, which determines the size of the groups of males where competition takes place, and *β*, which determines the probability of the highest ranked male in each group 'winning'. Threshold indicates the condition threshold below which a male will follow alternative mating tactics. Higher values for the critical rate of environmental change of a population indicate that the population can persist when environmental change is more rapid. All simulations were run for 500 time steps and other parameter values were *K_cap* = 500, *max_offspring* = 6, *and ART_success* = 0.5.

#### Effects of increasing the success of minor males

Figure S2 shows the effect of increasing the success of minor males in acquiring mates via the ART. As can be seen, increasing the number of matings that these males achieve does reduce the critical rate of environmental change, whether the ART is fixed or simultaneous. Notably, however, even with substantial increases in the mating success of males pursuing the ART the critical rate remains higher than it would be in the absence of such males when the ART is fixed. Indicating that this effect of fixed ARTs is robust even when the males pursuing the alternative tactic are able to gain matings successfully.

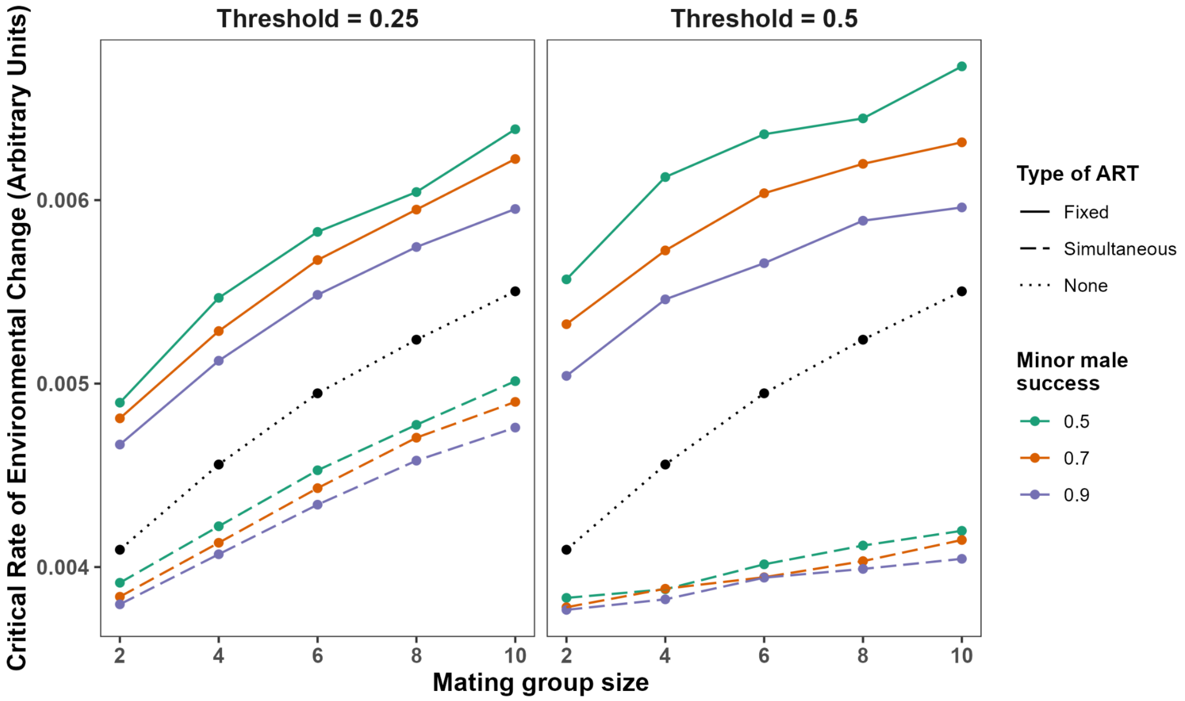


**Figure S2:** the effect of changing the probability of success for males pursuing alternative tactics on the critical rate of environmental change.variation in the critical rates of environmental change. “Minor male success” is the parameter *ART_success* from the simulation which determines the probability of a male pursuing the alternative tactic achieving a mating. Mating group size is the parameter *group_size*, which determines the size of the groups of males where competition takes place. Threshold indicates the condition threshold below which a male will follow alternative mating tactics. Higher values for the critical rate of environmental change of a population indicate that the population can persist when environmental change is more rapid. All simulations were run for 500 time steps and other parameter values were *K_cap* = 500, *max_offspring* = 6, *and* $\beta$ = 2.

Chevin L-M, Lande R, Mace GM. 2010. Adaptation, Plasticity, and Extinction in a Changing Environment: Towards a Predictive Theory. *PLoS Biol* **8**:e1000357.

Venables WN, Ripley BD. 2002. Modern Applied Statistics with S.
